# Nuclear import of *Arabidopsis* POLY(ADP-RIBOSE) POLYMERASE 2 is mediated by importin-α and a nuclear localization sequence located between the predicted SAP domains

**DOI:** 10.1101/304527

**Authors:** Chao Chen, Ruth Lintermann, Lennart Wirthmueller

## Abstract

Proteins of the poly(ADP-ribose) polymerase (PARP) family modify target proteins by covalent attachment of ADP-ribose moieties onto amino acid side chains. In *Arabidopsis*, PARP proteins contribute to repair of DNA lesions and modulate plant responses to various abiotic and biotic stressors. *Arabidopsis* PARP1 and PARP2 are nuclear proteins and given that their molecular weights exceed the diffusion limit of nuclear pore complexes, an active import mechanism into the nucleus is likely. Here we use confocal microscopy of fluorescent protein-tagged *Arabidopsis* PARP2 and PARP2 deletion constructs in combination with site-directed mutagenesis to identify a nuclear localization sequence in PARP2 that is required for nuclear import. We report that in co-immunoprecipitation assays PARP2 interacts with several isoforms of the importin-α group of nuclear transport adapters and that PARP2 binding to IMPORTIN-α2 is mediated by the identified nuclear localization sequence. Our results demonstrate that PARP2 is a cargo protein of the canonical importin-α/β nuclear import pathway.

## Introduction

The post-translational modification of proteins is an important component of plant responses to changes in their environment. Post-translational modifications (PTMs) are often rapid and reversible processes that allow plants to fine-tune the speed and duration of stress-induced signaling. Numerous PTMs have been described in plants including phosphorylation, ubiquitination, SUMOylation, acetylation and methylation (Hashiguchi and Komatsu, 2017; Withers and Dong, 2017). The N-terminal regions of the core histones are considered PTM hot spots and modification of histone tails plays a crucial role in transcriptional regulation and epigenetic processes including plant stress memory (Kleinmanns and Schubert, 2014; Asensi-Fabado et al., 2017).

Protein ADP-ribosylation is a PTM that recently has gained increasing attention in plants (Jia et al., 2013; Song et al., 2015; Feng et al., 2016; Vainonen et al., 2016; Rissel et al., 2017). Enzymes of the ADP-ribosyl transferase family catalyze covalent modification of proteins and DNA with ADP-ribose. In mammalian cells, ADP-ribosyl transferases play important roles in several cellular pathways including, amongst others, DNA damage repair, apoptosis, transcriptional regulation, mRNA stability and the cell cycle (Bai, 2015; Bock et al., 2015). Some ADP-ribosyl transferases catalyze the formation of poly-ADP-ribose chains by using the terminal ADP-ribose transferred onto an acceptor molecule as a substrate for subsequent rounds of ADP-ribosylation. These ADP-ribosyl transferases are also called <u>P</u>oly(<u>A</u>DP-<u>R</u>ibose) <u>P</u>olymerases (PARPs). Other mammalian ADP-ribosyl transferases only attach a single ADP-ribose moiety onto acceptor molecules and therefore act as mono-ADP-ribosyl transferases (Bock and Chang, 2016). ADP-ribosyl transferases use NAD^+^ as a co-substrate to transfer the ADP-ribose moiety from NAD^+^ onto an amino acid side chain. Similar to protein kinases, several ADP-ribosyl transferases not only ADP-ribosylate other proteins but also undergo auto-modification (Adamietz, 1987; Muthurajan et al., 2014; Vyas et al., 2014).

Protein ADP-ribosylation remains a poorly characterized PTM in plants. Although enzymes of the ADP-ribosyl transferase family are conserved in plants (Lamb et al., 2012; Vainonen et al., 2016), only few ADP-ribosylated acceptor proteins have been characterized (Feng et al., 2015, 2016). In addition, the consequences of protein ADP-ribosylation in plants remain largely unknown. Notably, the best-characterized examples of plant protein modification by ADP-ribosylation come from studies of plant pathogens that transfer effector proteins with ADP-ribosyl transferase activity into plant cells to suppress activation of plant innate immunity (Singer et al., 2004; Fu et al., 2007; Wang et al., 2010; Jeong et al., 2011; Nicaise et al., 2013).

The *Arabidopsis thaliana* genome encodes three PARP proteins (annotated as PARP1-3) with 27-47% sequence identity to *Hs*PARP-1 and *Hs*PARP-2. *Arabidopsis* PARP1 and PARP2 are active enzymes while direct evidence for PARP3 ADP-ribosyl transferase activity is lacking (Babiychuk et al., 1998; Feng et al., 2015). Treatment of *Arabidopsis* with γ-radiation or genotoxic agents activates PARP1 and PARP2 (Song et al., 2015). Based on the analysis of *parp* single and double mutants, PARP1 and PARP2 fulfill partially redundant functions in response to genotoxic stress. Upon treatment of *Arabidopsis* seedlings with the DNA double strand break-inducing agent Bleomycin, PARP2 mediates the majority of detectable poly-ADP-ribosylation (Song et al., 2015). However, based on quantification of DNA damage via the Comet assay, *parp1* mutants show higher levels of DNA damage compared to *parp2* mutants following exposure to methyl methane sulfonate as well as in untreated seedlings (Jia et al., 2013). How DNA damage enhances the enzymatic activity of plant PARPs has not been reported in detail. However, based on sequence conservation between plant PARPs and mammalian homologs, the access of NAD^+^ to the active site might be blocked by a protein regulatory domain (PRD) located N-terminal to the catalytic domain. Based on the analysis of human PARP-1, sensing of DNA double strand breaks by the N-terminal domains might result in a conformational change of the PRD thereby relieving auto-inhibition of the catalytic domain (Langelier et al., 2012, 2018). In human PARP-1, the binding site for DNA double strand breaks is formed by two Zinc finger domains and a WGR domain (conserved Trp, Gly and Arg residues) (Langelier et al., 2012). Similar to human PARP-1, predicted Zn finger and WGR domains appear to be conserved in *Arabidopsis* PARP1. In contrast, for *Arabidopsis* PARP2, two N-terminal SAP (<u>S</u>AF-A/B, <u>A</u>cinus and <u>P</u>IAS) domains followed by a WGR domain have been predicted suggesting that the mechanism of PARP2 activation by DNA damage differs from PARP1 (Lamb et al., 2012; Vainonen et al., 2016).

PARP1 and PARP2 localize to the plant cell nucleus consistent with their roles in DNA damage repair (Babiychuk et al., 2001; Song et al., 2015). Given their entirely nuclear localization and predicted molecular weights of 111 (PARP1) and 72 (PARP2) kDa, an active nuclear import mechanism for plant PARPs is likely. For PARP2 Babiychuck *et al.* (2001) reported that a GFP fusion of an N-terminal fragment spanning amino acids 1-104 is entirely nuclear localized, indicating an active import mechanism. Active transport processes through nuclear pore complexes are mediated by several distinct transport systems. Karyopherins of the importin-α/β group function as adapter proteins that bind cargoes with exposed nuclear localization sequences (NLS) in the cytoplasm, transport them through nuclear pore complexes and release their cargoes in the nucleoplasm (Christie et al., 2016). While importin-β proteins can achieve active transport across the nuclear envelope by directly interacting with Phe/Gly-rich repeats of nucleoporins that line the nuclear pore channel (Allen et al., 2001; Bayliss et al., 2002), importin-α proteins form a ternary complex with their cargoes and importin-β for transport to the nucleus (Weis et al., 1996; Cingolani et al., 1999). In *Arabidopsis*, there are nine importin-α isoforms that show both redundant but also specific transport functions (Huang et al., 2010; Merkle, 2011; Wirthmueller et al., 2013; Roth et al., 2017). While the NLS that mediate binding to importin-β can diverge in sequence (Christie et al., 2016), importin-α proteins bind canonical mono- and bipartite NLS characterized by Lys/Arg-rich consensus sequences (Marfori et al., 2011, 2012). Here we identify an NLS in PARP2 that is sufficient and required for nuclear import. We further demonstrate that this NLS mediates binding to *Arabidopsis* IMPORTIN-α2 in plant cell extracts.

## Materials and Methods

### Plants and growth conditions

*Nicotiana benthamiana* plants were grown in a green house at 22 °C/20 °C day/night temperatures and 16 h (06:00 to 22:00) supplemental light (200-230 μmol m^2^ s^-1^) from tungsten lamps.

### Generation of plant and *E. coli* expression constructs

The following Gateway-compatible pENTR4 plasmids were generated for this work: pENTR4-PARP2, pENTR4-PARP2^SAP-WGR^, pENTR4-PARP2^PRD-CAT^, pENTR4-PARP2^SAP^, pENTR4-PARP2^WGR^, pENTR4-PARP2^48-51AAAA^, pENTR4-PARP2^48-51QMQL^, pENTR4-PARP2^SAP48-51AAAA^, pENTR4-PARP2^SAP48-51QMQL^, and pENTR4-PARP2^SAP92/93AA^. All pENTR4 constructs lack a stop codon for translational fusion to xFP reporters. PARP2 mutant constructs were generated either by the QuikChange method (pENTR4-PARP2^48-51AAAA^ and pENTR4-PARP2^SAP92/93AA^) or by splice-by-overlap-extension (SOE) PCR (pENTR4-PARP2^48-51QMQL^). To generate translational fusions between PARP2 variants and xFP tags, Gateway LR reactions between PARP2 (or PARP2 deletion) constructs and pK7FWG2 (enhanced GFP tag) or pH7RWG2 (RFP tag) (Karimi et al., 2002) were performed. The IMPORTIN-α6:GFP construct was created by an LR reaction between pENTR/D-Topo IMPORTIN-α6 (Roth et al., 2017) and pK7FWG2. The other GFP-tagged importin-α plant expression constructs have been described previously (Wirthmueller et al., 2015).

To generate the *E.coli* expression construct for ΔIBB IMPORTIN-α2 [lacking the auto-inhibitory importin-β-binding (IBB) domain], a cDNA fragment coding for amino acids 75-535 was amplified and cloned into KpnI/HindIII-linearized pOPINF (Berrow et al., 2007) via Gibson assembly. The *E. coli* PARP2 SAP (amino acids 1-105) expression construct was generated following the same strategy but using pOPINS3C (Bird, 2011) as expression plasmid. For production of α-GFP affinity beads, the coding sequence of an α-GFP nanobody (Addgene plasmid #49172; Kubala et al., 2010) was fused in frame with a Gly-Gly-Ser-Gly-Ser linker and the Halo tag from plasmid pGW-nHalo (Peterson and Kwon, 2012) into pOPINE (Berrow et al., 2007) via Gibson assembly. See Table S1 for oligo nucleotides and cloning methods.

### Transient protein expression in *N. benthamiana*

For transient expression in *N. benthamiana* leaves binary vectors were transformed into *Agrobacterium tumefaciens* strain GV3101::pMP90. Agrobacteria were plated on YEB medium with appropriate antibiotics and incubated for 3 days at 28 °C. On the day of infiltration, the cells were resuspended in infiltration medium (10 mM MES pH 5.6, 10 mM MgCl_2_) and the OD_600_ was adjusted to 0.8. To suppress transgene silencing Agrobacteria expressing the tomato bushy stunt virus 19K silencing suppressor were coinfiltrated. The culture of the 19K strain was adjusted to an OD_600_ of 6.0. After adding Acetosyringone to a final concentration of 100 μM the bacteria were incubated for 2 h at RT and then mixed in a ratio of xFP[20]:19K[3] for localization experiments or GFP[10]:RFP[10]:19K[3] for co-immunoprecipitation experiments. *Agrobacterium* mixtures were infiltrated with a needleless syringe into leaves of 4-5 week-old *N. benthamiana* plants and leaf material was harvested for microscopy or protein extraction 72 h later.

### Microscopy

Leaf discs excised from *N. benthamiana* were mounted on microscope slides in water and the subcellular localization of xFP-tagged proteins was analyzed using a Leica TCS SP5 confocal unit attached to a Leica DMI6000 CS microscope. GFP and YFP were excited at 488 nm and collected at 500-525 nm and 525-540 nm, respectively. RFP was excited at 561 nm and collected at 580-610 nm. The gain setting of the confocal unit was adjusted to just below the threshold for saturation of the signal. Images were acquired and analyzed using LAS AF software (Leica).

### Plant protein extraction, immunoprecipitation and detection

Protein extracts were prepared by grinding *N. benthamiana* leaf material in liquid nitrogen to a fine powder followed by resuspension in extraction buffer [50 mM Tris, 150 mM NaCl, 10% Glycerol, 1 mM EDTA, 5 mM DTT, 1x Protease inhibitor cocktail (Sigma, http://www.sigmaaldrich.com), pH 7.5] at a ratio of 2 ml buffer per 1 g leaf material. The extracts were centrifuged at 20000 x *g* / 4°C / 20 min and the supernatant was either boiled in Sodium dodecyl sulfate (SDS) sample buffer for western blots or used for co-immunoprecipitation experiments. For western blots protein samples were separated by SDS-PAGE and electro-blotted onto nitrocellulose membrane. Antibodies used were α-GFP 210-PS-1GP (Amsbio, http://www.amsbio.com) and α-mCherry ab125096 (Abcam, http://www.abcam.com). For co-immunoprecipitation a fraction of the supernatant was saved as ‘input’ sample and 15 μl of α-GFP-nanobody:Halo:His6 magnetic beads (see below) were added to 1.4 ml of the remaining supernatant. The samples were incubated on a rotating wheel at 4°C for 2 h followed by collection of the beads using a magnetic sample tube rack. The beads were washed 3 times with 1 ml extraction buffer and then boiled in 40 μl SDS sample buffer to elute protein from the beads.

### Protein expression and purification

For protein expression in *E. coli*, pOPINS3C carrying His6:SUMO:SAP and pOPINE carrying the α-GFP-nanobody:Halo:His6 construct were transformed into SHuffle^®^ T7 Competent *E. coli* cells (New England Biolabs). The His6:ΔIBB-IMPORTIN-α2 protein was expressed from pOPINF in SoluBL21 cells (Genlantis). All cultures were grown to an OD_600_ of 1 to 1.2 at 37 °C, then shifted to 18 °C and expression was induced by addition of 0.5 mM IPTG. After 16-18 h the bacteria were harvested by centrifugation (5000 x *g*, 12 min.). His6-tagged proteins were purified using a combination of immobilized metal ion affinity chromatography and size exclusion chromatography. To this end, the cell pellets were resuspended in 50 mM Tris, 300 mM NaCl, 50 mM Glycine, 20 mM Imidazole, 5% Glycerol, pH 8.0 at a ratio of 20 ml per initial litre of culture volume. Bacterial lysis was achieved by incubation with Lysozyme (20 min., RT) followed by sonication (2 times 2 min. at level 4 on a Branson Sonifier 150). The cell extract was cleared from debris and insoluble proteins by centrifugation (30000 x *g*, 4 ºC, 20 min) and the supernatant was loaded onto pre-equilibrated Ni^2+^-immobilized metal ion affinity chromatography columns (His-Trap HP 5 ml, GE Healthcare). After washing out unbound proteins, His6-tagged proteins were eluted with 50 mM Tris, 300 mM NaCl, 50 mM Glycine, 250 mM Imidazole, 5% Glycerol, pH 8.0 and directly injected onto Hi-Load 26/60 Superdex 75 (His6:SUMO:SAP, α-GFP-nanobody:Halo:His6) or Superdex 200 (His6:ΔIBB-IMPORTIN-α2) size exclusion columns (GE Healthcare) using buffer A4 (20 mM HEPES, 150 mM NaCl, pH 7.5) as the elution buffer. Proteins were concentrated using ultrafiltration columns (Sartorius), frozen in liquid nitrogen and stored at −80 ºC.

### Analytical size exclusion chromatography

Purified His6:ΔIBB-IMPORTIN-α2 or His6:SUMO:SAP was injected onto a Superdex 200 10/300 GL size exclusion column (GE Healthcare) in a volume of 1 ml. Proteins were eluted from the column with buffer A4 at a flow rate of 0.4 ml/min and 1 ml fractions were collected. These fractions were subsequently analyzed by SDS-PAGE and Instant Blue (Expedeon) staining.

### Production of GFP affinity magnetic beads

To couple the α-GFP nanobody:Halo:His6 fusion protein to magnetic beads, 250 μl of Magne^®^ HaloTag^®^ Beads (Promega) were washed in 1 ml A4 buffer and the beads were collected using a magnetic sample tube rack. 1.5 mg of the α-GFP nanobody:Halo:His6 protein was mixed with the washed beads in 2 ml buffer A4 and incubated on a rotating wheel for 2 h at 4 °C and 15 rpm. The magnetic beads were collected and washed three times with 1 ml A4 buffer. The affinity beads were stored in 250 μl of storage buffer (50 mM HEPES, 150 mM NaCl, 15% Glycerol, 0.05% NaNs, pH 7.5) at 4 °C until use.

### Accession numbers

*Arabidopsis thaliana* PARP2 (AT4G02390), IMPORTIN-α1 (AT3G06720), IMPORTIN-α2 (AT4G16143), IMPORTIN-α3 / MOS6 (AT4G02150), IMPORTIN-α4 (AT1G09270), IMPORTIN-α6 (AT1G02690), IMPORTIN-α9 (AT5G03070); *Hyaloperonospora arabidopsidis* effector protein HaRxL106 (GenBank HE574762.1).

## Results

### PARP2 interacts with several importin-α proteins

We expressed PARP2:RFP in epidermal *N. benthamiana* cells and confirmed that the fusion protein localized to the nucleus (Fig. 1a). In contrast, free RFP showed a nucleo-cytoplasmic distribution, which is in accordance with unrestricted passive diffusion of macromolecules of molecular weights below 40-60 kDa through nuclear pore complexes (Timney et al., 2016). To test for interaction with importin-α, we co-expressed PARP2:RFP with the six importin-α proteins that are expressed in *Arabidopsis* leaf tissue (IMPORTIN-α1-4, IMPORTIN-α6, IMPORTIN-α9; Wirthmueller et al., 2013). For immunoprecipitation, and based on the finding that a C-terminal GFP tag does not interfere with IMPORTIN-α3 function (Palma et al., 2005), we expressed all importin-α proteins as N-terminal fusions to GFP. As shown in Fig. 1b, PARP2:RFP coprecipitated with several importin-α isoforms. In three independent experiments we observed a trend for stronger interaction between PARP2:RFP and IMPORTIN-α2 and IMPORTIN-α4 compared to the other importin-α proteins (Fig. 1b; Fig. S1). PARP2:RFP showed no interaction with YFP that we used as negative control (Fig. 1b; Fig. S1). Compared to the previously characterized interaction between IMPORTIN-α2 and the oomycete effector protein HaRxL106 (Wirthmueller et al., 2015), PARP2:RFP showed relatively weak binding to importin-α proteins (Fig. 1b; Fig. S1).

**Figure 1.**
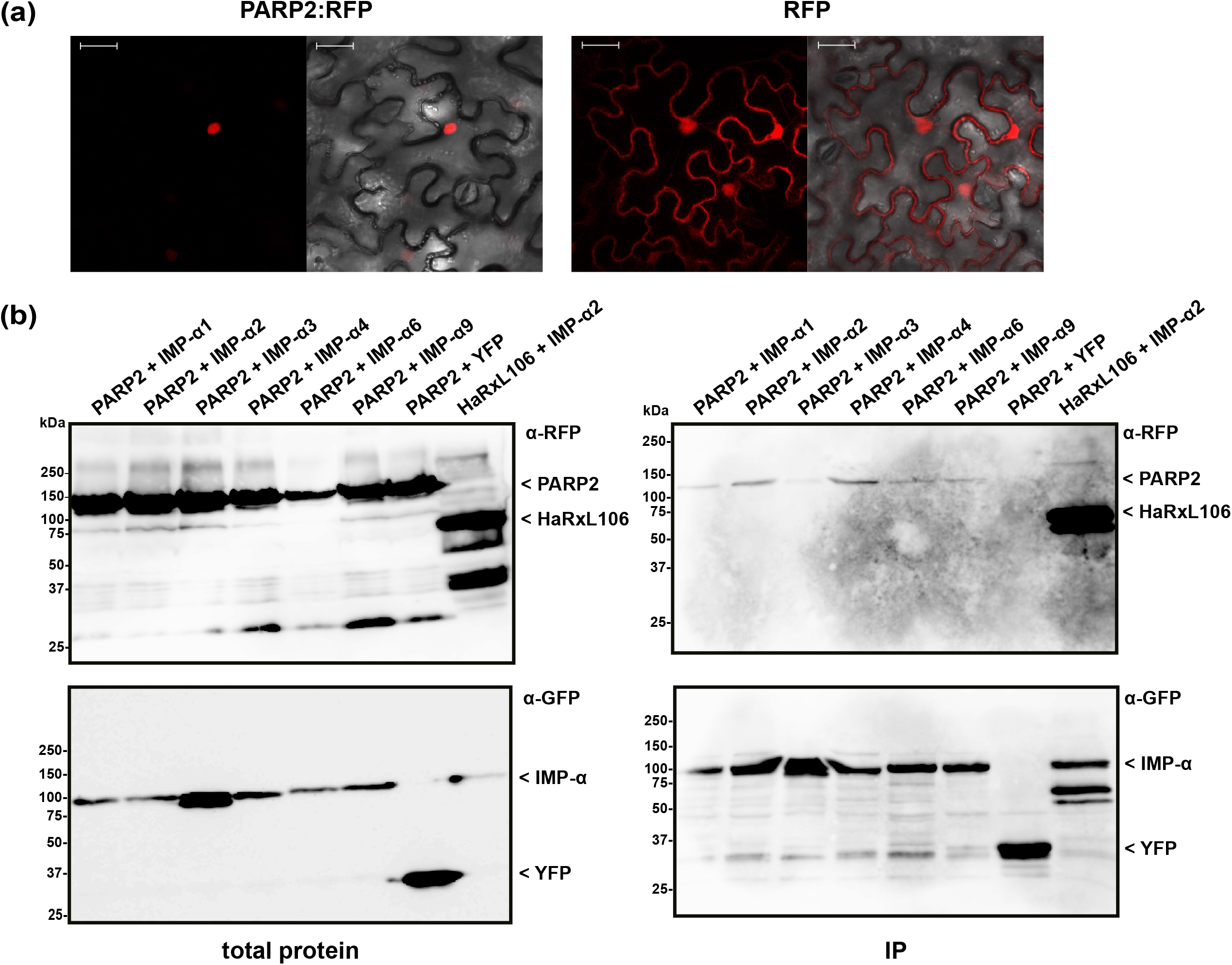
PARP2 localizes to the nucleus and interacts with several importin-α isoforms. **(a)** Representative (n = 10) confocal microscopy images of *N. benthamiana* cells expressing PARP2:RFP or free RFP. Left image RFP channel. Right image overlay with bright field image. Scale bar 30 μm. Images were taken 72 h after infiltration. **(b)** GFP-tagged versions of IMPORTIN-α1, - α2, - α3, - α4, - α6, - α9 or free YFP were coexpressed with PARP2:RFP or RFP:HaRxL106 (positive control) in *N. benthamiana*. At 72 h post infiltration, GFP-tagged proteins or YFP were immunoprecipitated and co-precipitating PARP2:RFP was detected by an α-RFP western blot.

### A nuclear localization sequence located between the PARP2 SAP domains is sufficient and required for nuclear import

To map sequences of PARP2 that are required for nuclear import we generated truncated variants of the protein and expressed these as GFP fusions in *N. benthamiana*. Based on the predicted domain structure of PARP2 (Fig. 2a), we initially split the protein between the WGR domain and the protein regulatory domain (PRD). The fusion of the SAP-WGR domains to GFP retained an entirely nuclear localization (Fig. 2b). In contrast, a GFP fusion of the PARP2 fragment comprising PRD and catalytic (CAT) domains showed a nucleo-cytoplasmic distribution (Fig. 2b). Therefore, the sequence(s) for active nuclear import of PARP2 are located within the first 280 amino acids of the protein and the regulatory and catalytic domains do not contain additional NLS. We then expressed the isolated SAP and WGR domains as GFP fusions in *N. benthamiana*. The SAP:GFP fusion showed an entirely nuclear localization while the WGR:GFP signal was distributed between the cytoplasm and the nucleus (Fig. 2b). This suggests that all nuclear targeting sequences of PARP2 are located within the first 127 amino acids of the protein that are predicted to fold into two SAP domains (Lamb et al., 2012; Vainonen et al., 2016).

**Figure 2.**
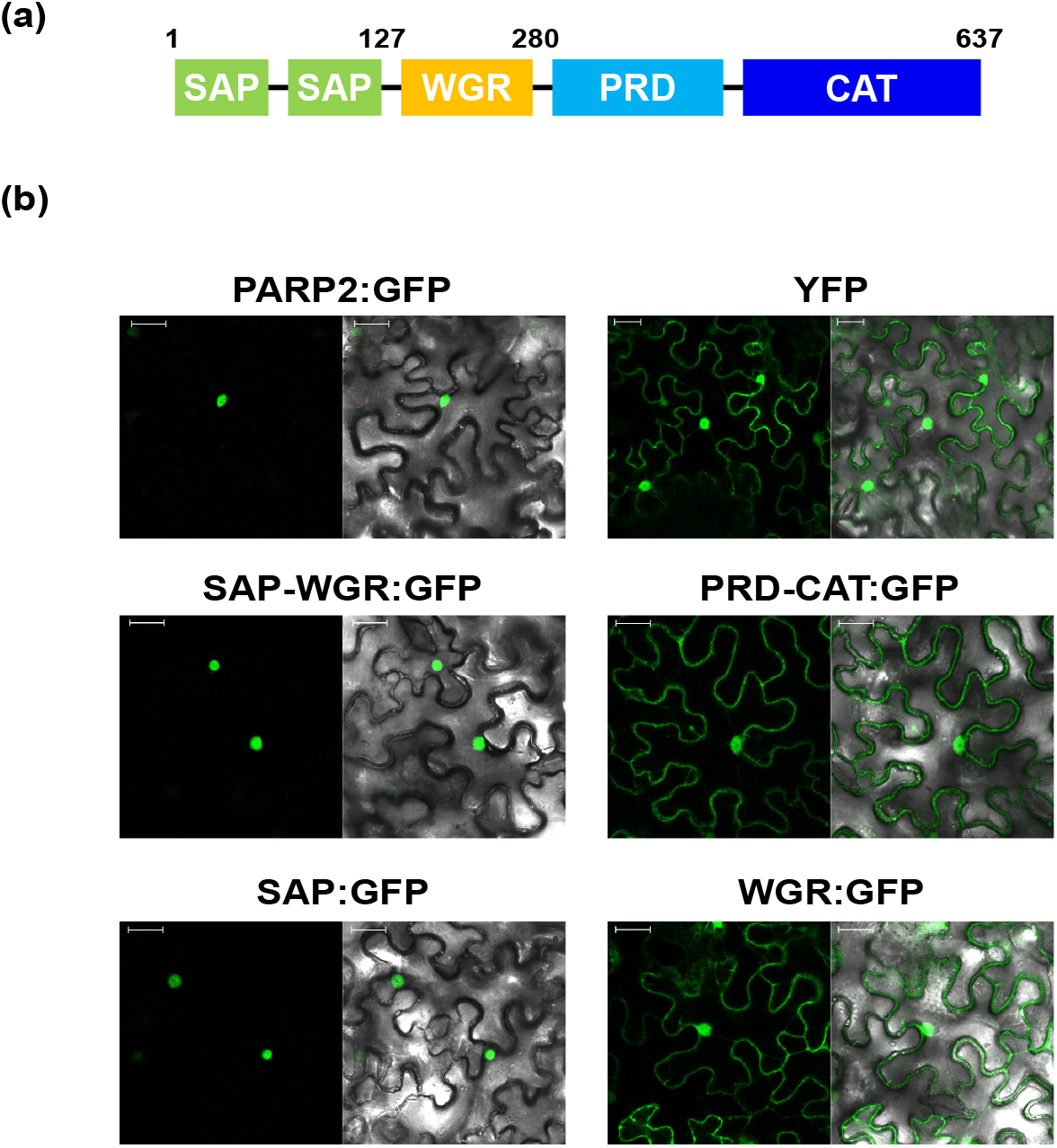
PARP2 nuclear targeting sequences are located in the N-terminal SAP domains. **(a)** Predicted domain structure of PARP2 based on homology modeling and sequence conservation. SAP = <u>S</u>AF-A/B, <u>A</u>cinus and <u>P</u>IAS domains; WGR = domain with conserved Trp, Gly and Arg residues; PRD = protein regulatory domain; CAT = catalytic domain. **(b)** Representative (n = 10) confocal microscopy images of *N. benthamiana* cells expressing the indicated GFP fusion proteins or free YFP. Left image GFP or YFP channel. Right image overlay with bright field image. Scale bar 30 μm. Images were taken 72 h after infiltration.

Having mapped the NLS of PARP2 to the N-terminal 127 amino acids, we focused on clusters of basic amino acids that could constitute an NLS. Using NLS Mapper (Kosugi et al., 2009), we identified a candidate monopartite NLS (SKSKRKRNS; amino acids 45-53; NLS Mapper score 7.0) that might be part of a larger bipartite NLS (KSKRKRNSSNDTYESNKLIAI, amino acids 46-66; NLS Mapper score 6.5). We mutated the KRKR motif of the candidate NLS either to a quadruple Alanine or the sequence QMQL. The latter sequence is more similar to KRKR with respect to the length of amino acid side chains but does not carry a basic charge. When we transiently expressed the corresponding mutant SAP:RFP fusions in *N. benthamiana* we observed a nucleo-cytoplasmic distribution of the RFP signal, indicating that PARP2 amino acids 48-51 are indeed part of an NLS (Fig. 3a). In contrast, changing two other basic amino acids of PARP2 (K92/K93) to Alanine did not result in a nucleo-cytoplasmic distribution (Fig. 3a). We then introduced the AAAA and QMQL NLS mutations into full-length PARP2 and analyzed the subcellular localization of the respective fusion proteins. As shown in Fig. 3a, both mutations in the NLS resulted in a predominantly cytoplasmic localization of PARP2:RFP with strong depletion of the signal from the nucleus. In cases where we observed a residual RFP signal from the nucleus the fluorescence was stronger at the nuclear envelope (Fig. 3b). Therefore amino acids 48-51 are essential for nuclear import of PARP2 and, based on the cytoplasmic localization of the NLS mutant variants, it appears unlikely that PARP2 two has other strong nuclear targeting sequences.

**Figure 3.**
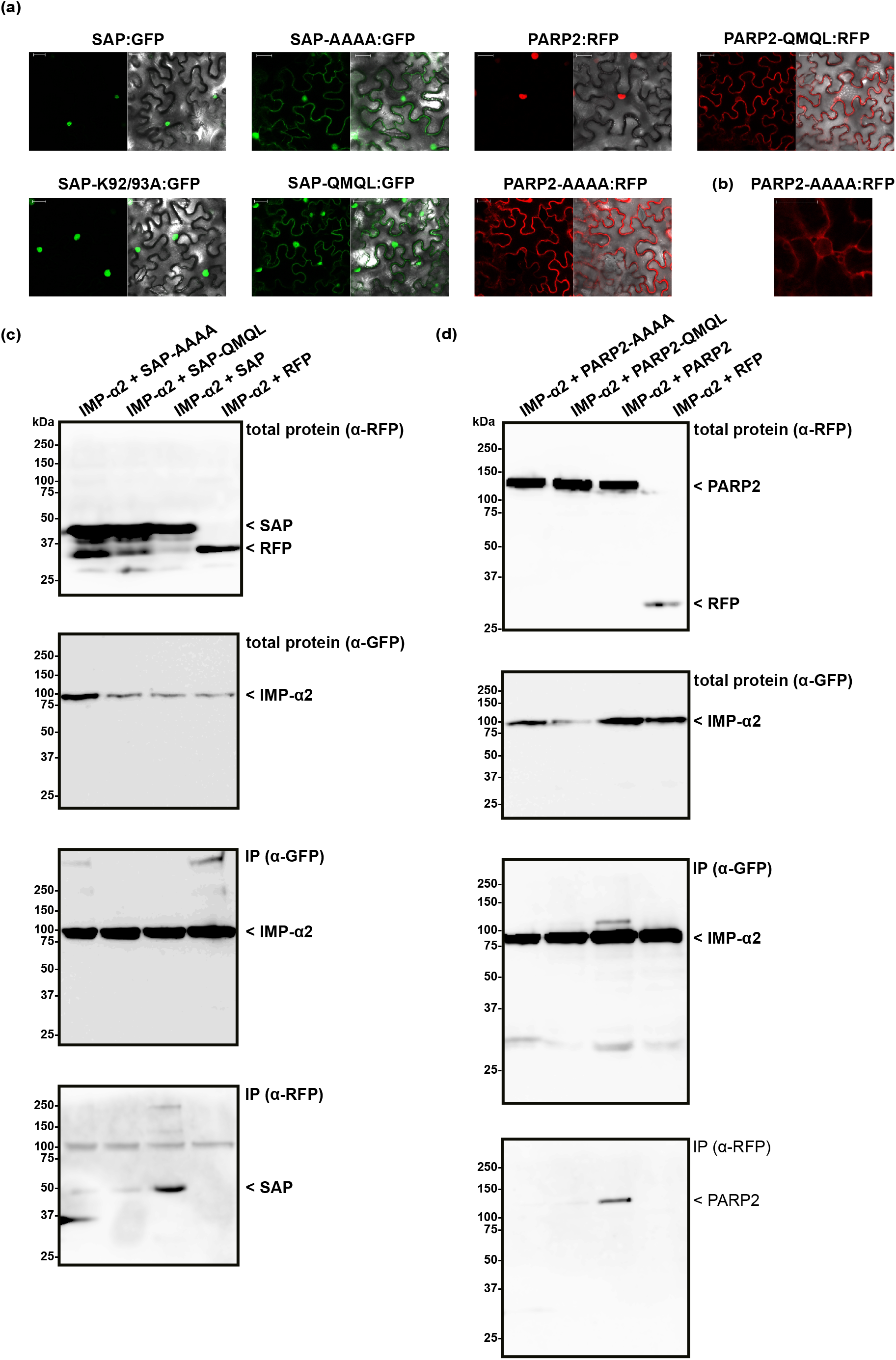
PARP2 amino acids 48-51 are essential for nuclear targeting and mediate interaction with IMPORTIN-α2. **(a)** Representative (n = 10) confocal microscopy images of *N. benthamiana* cells expressing the indicated GFP/RFP fusion proteins. Left image GFP or RFP channel. Right image overlay with bright field image. Scale bar 30 μm. Images were taken 72 h after infiltration. ‘AAAA’ indicates mutation of PARP2 amino acids 48-51 to quadruple Ala. ‘QMQL’ indicates mutation of PARP2 amino acids 48-51 to Gln-Met-Gln-Leu. ‘K92/93A’ indicates mutation of PARP2 Lys 92 and 93 to Ala. **(b)** Distribution of the RFP signal from PARP2-AAAA:RFP around a nucleus (n = 4). **(c)** IMPORTIN-α2:GFP was co-expressed with free RFP or the indicated RFP-tagged variants of the PARP2 SAP domains in *N. benthamiana*. At 72 h post infiltration, IMPORTIN-α2:GFP was immunoprecipitated and co-precipitating RFP-tagged proteins were detected by an α-RFP western blot. **(d)** Co-immunoprecipitation experiment as in (c) but with the wildtype and mutated full-length PARP2 sequences instead of the SAP domains.

To test whether nuclear import of PARP2 correlates with binding to importin-α, we performed co-immunoprecipitation experiments between IMPORTIN-α2 and mutated variants of the isolated SAP domains and full-length PARP2. As shown in Fig. 3c, mutation of the KRKR motif resulted in weaker or non-detectable binding of the SAP domains to IMPORTIN-α2 (see Fig. S2 for data from two additional independent experiments). Likewise, PARP2 variants with mutations of the KRKR motif showed quantitatively reduced or no detectable binding to IMPORTIN-α2 (Fig. 3d; Fig. S3). Overall, we observed a correlation between the strength of IMPORTIN-α2 binding and nuclear localization for the isolated SAP domains and full-length PARP2. These results are consistent with importin-α-dependent nuclear import of PARP2 mediated by the NLS comprising the KRKR motif.

### The PARP2 SAP domains and IMPORTIN-α2 do not form a stable complex *in vitro*

Several NLS bind to importin-α with affinities in or below the low micro-molar range (Hübner et al., 1999; Hodel et al., 2006; Timney et al., 2006; Kosugi et al., 2008; Chang et al., 2012). This often allows for isolation of a stable protein complex between NLS peptides and the Armadillo repeat domains of importin-α proteins (Conti et al., 1998; Fontes et al., 2003; Marfori et al., 2012). In comparison to the interaction between the oomycete effector HaRxL106 and IMPORTIN-α3, for which a stable complex could be detected (Wirthmueller et al., 2015), PARP2 showed weaker binding to all tested importin-α isoforms (Fig. 1b). To test for protein complex formation between the SAP domains of PARP2 and IMPORTIN-α2, we expressed the SAP domains (amino acids 1-105) and the Armadillo repeat domain of IMPORTIN-α2 (amino acids 75-535) as His6-tagged proteins in *E. coli* (for the SAP domains we produced a His6:SUMO:SAP fusion, see methods for details). We purified both proteins by immobilized metal ion affinity and subsequent size exclusion chromatography, mixed the proteins in a 1:2 (IMPORTIN-α2:SAP) molar ratio and assessed complex formation by analytical size exclusion chromatography. As shown in Fig. 4a, the IMPORTIN-α2 and SAP proteins eluted in separate peaks when injected individually onto the size exclusion column. When we mixed the two proteins we still detected two separate peaks and no higher molecular weight peak. Analyzing the eluted fraction by SDS-PAGE showed that the elution profile of IMPORTIN-α2 was not altered by the 1-fold molar excess of the PARP2 SAP domains (Fig. 4b; Fig. S4). This suggests that unlike other importin-α/NLS pairs the PARP2 SAP domains do not form a stable complex with IMPORTIN-α2 under the conditions tested here. As we have not been successful in expressing the SAP domains with a shorter affinity tag, we could not assess whether the His6:SUMO tag N-terminal to the PARP2 SAP domains might interfere with IMPORTIN-α2 binding.

**Figure 4.**
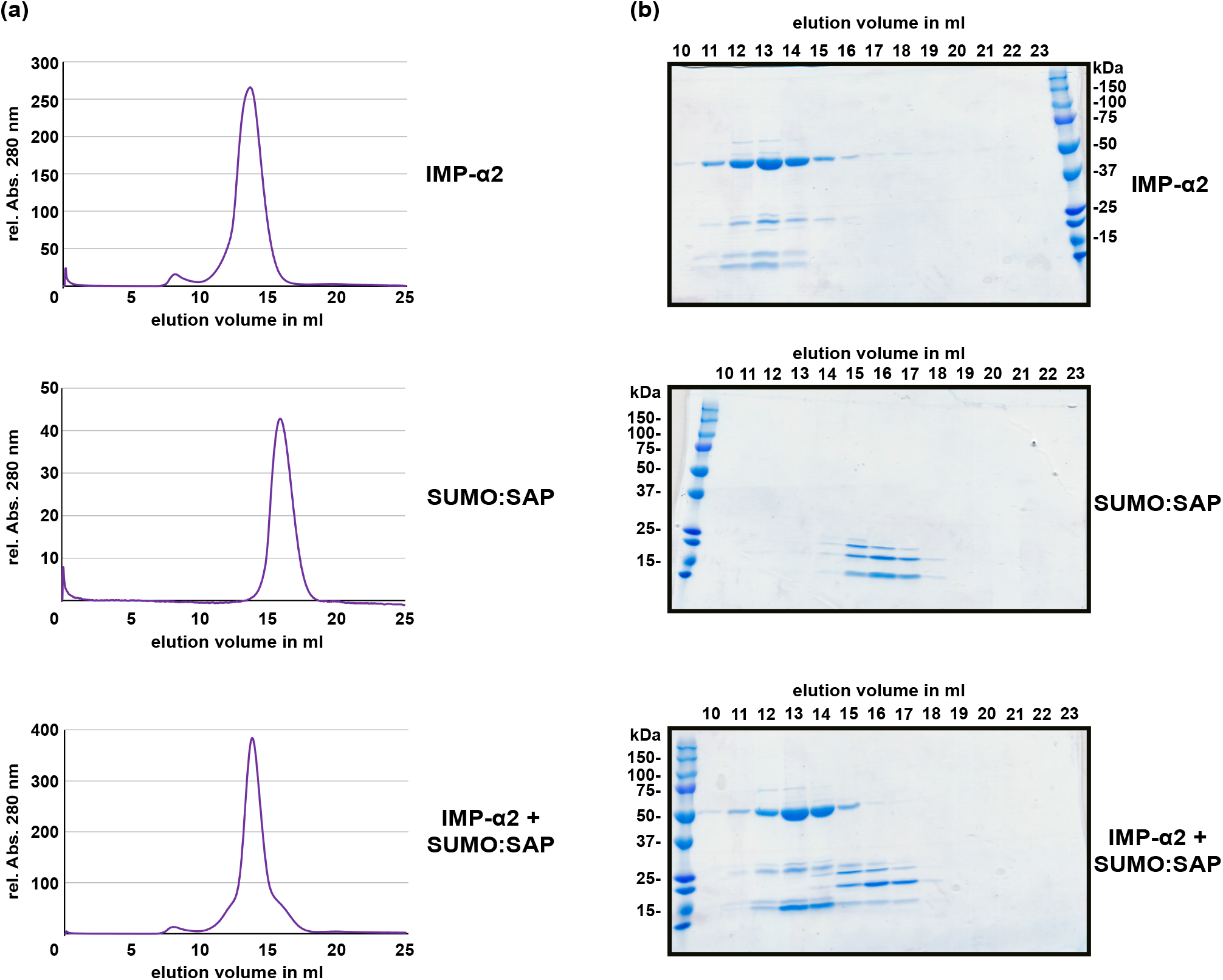
The PARP2 SAP domains and IMPORTIN-α2 do not form a stable protein complex *in vitro*. **(a)** Size exclusion chromatography elution profiles of His6:ΔIBB-IMPORTIN-α2, His6:SUMO:SAP and a mixture of both proteins (1:2 molar ratio) as determined by the absorption at 280 nm. **(b)** Coomassie-stained SDS-PAGE of the elution fractions from (a). The molecular weight of His6:ΔIBB-IMPORTIN-α2 is 53 kDa. The molecular weight of the His6:SUMO:SAP fusion is 25 kDa.

## Discussion

We present evidence for an active importin-α-mediated nuclear import of *Arabidopsis* PARP2. This model is supported by i) the identification of an NLS in PARP2 that is sufficient and required for nuclear import and ii) binding studies showing that this NLS mediates interaction with IMPORTIN-α2 in plant cells (Figs. 2 and 3). PARP2 preferentially bound to IMPORTIN-α2 and IMPORTIN-α4 although we detected weaker interactions with other importin-α isoforms (Fig. 1b; Fig. S1). This is consistent with partially redundant functions of importin-α isoforms in nuclear transport although examples of specific importin-α/cargo interactions have also been reported (Palma et al., 2005; Timney et al., 2006; Bhattacharjee et al., 2008; Wirthmueller et al., 2015; Roth et al., 2017).

While in some experiments mutation of the KRKR motif in PARP2 completely abolished binding to IMPORTIN-α2, we observed a quantitative reduction of the interaction in other experiments (Figs. 3c and 3d; Figs. S2 and S3). Based on the NLS Mapper prediction it is possible that the KRKR motif is part of a larger bipartite NLS. Two clusters of basic amino acids that make contact to the major and minor NLS binding sites of importin-α, respectively, characterize canonical bipartite NLS (Marfori et al., 2011). Therefore, the weak association between NLS mutant variants of the SAP domains or full-length PARP2 and IMPORTIN-α2 that we observed in some experiments could be due to other basic amino acids that contribute to importin-α binding. However, the cytoplasmic localization of PARP2 NLS mutant variants (Figs. 3a) suggests that if such a second contact point between PARP2 and importin-α exists it is not sufficient for nuclear import.

The NLS that we identified here is located between the two predicted SAP domains of PARP2. The location of the NLS in a short stretch of amino acids with no predicted secondary structure is consistent with the finding that NLS are often located in disordered regions of proteins. The identified NLS at position 48-51 of PARP2 is consistent with data from Babiychuck et al. (2001) who reported that the first 104 amino acids of the protein are sufficient for nuclear import. Based on an alignment of PARP2 sequences from other plant species (Fig. 5), the KRKR motif is partially conserved in PARP2 homologs. In several of the sequences shown in Fig. 5, amino acids with polar side chains or Gly are present at the position corresponding to *Arabidopsis* PARP2 K48. However, in all of these cases an Arg is present five amino acids downstream of the polar amino acid/Gly suggesting that in these sequences the cluster of basic amino acids is slightly shifted. Therefore, a cluster of basic amino acids that could mediate binding to importin-α is conserved in PARP2 homologs. Notably, also the nuclear targeting signal for mouse PARP2 is located in its N-terminal DNA-binding domain and two Lysine residues that are critical for nuclear import are modified by acetylation (Haenni et al., 2008a, 2008b).

**Figure 5.**
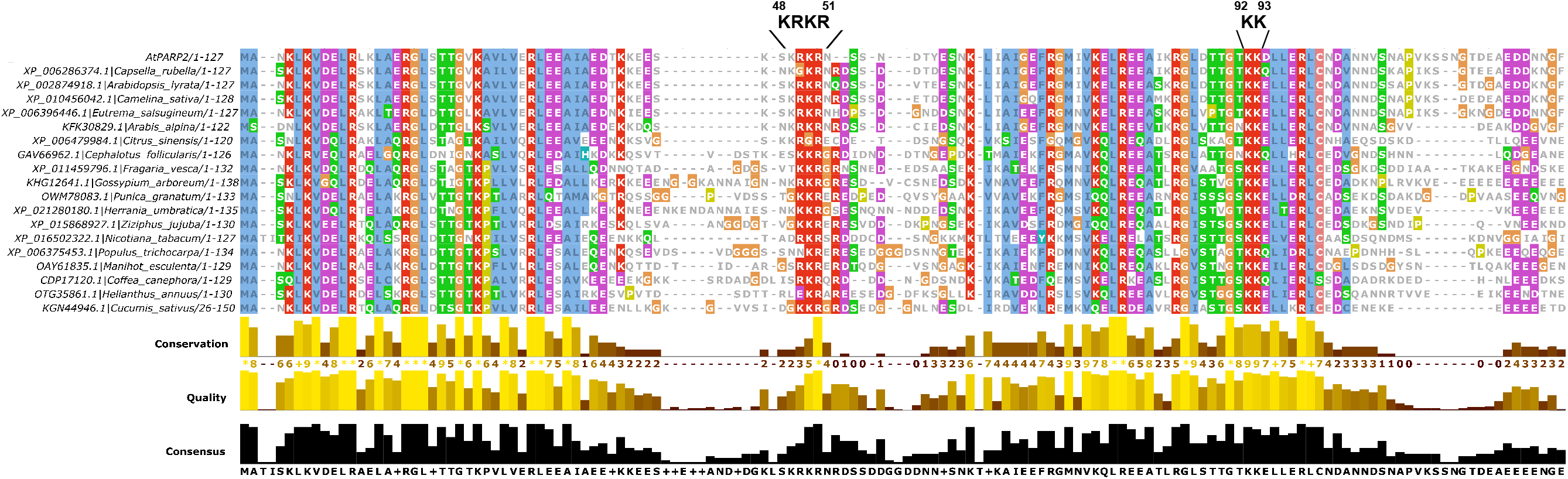
Multiple sequence alignment showing the partial conservation of the KRKR motif (PARP2 amino acids 48-51) in PARP2 homologs from different plant species. Only the SAP domains are shown. The sequence alignment was generated with Clustal Omega (Sievers and Higgins, 2014).

We noticed that in direct comparison to the interaction between IMPORTIN-α2 and the oomycete effector protein HaRxL106, PARP2 showed a comparably weak association with IMPORTIN-α2 as well as other importin-α isoforms (Figs. 1b). This is also consistent with no detectable interaction between the SAP domains and IMPORTIN-α2 by size exclusion chromatography (Fig. 4). The reported dissociation constants between NLS and importin-α proteins span a surprisingly large range, which might in part be explained by the different methods employed for their analyses (Marfori et al., 2012; Wirthmueller et al., 2015; de Barros et al., 2017). Although we observed comparably weak interaction between PARP2 and importin-α in plant cell extracts and no binding *in vitro*, the affinity between PARP2 and importin-α is sufficient for nuclear import in plant cells (Figs. 1a). Regarding the results with recombinantly expressed proteins, PTMs of amino acids within or adjacent to NLS can alter the affinity for importin-α and it is possible that such an unknown PTM is absent in the *E. coli* expression system (Moll et al., 1991; Harreman et al., 2004; Róna et al., 2013). Alternatively, the His6:SUMO affinity tag might hinder binding of IMPORTIN-α2 to the PARP2 SAP domains.

In summary, our data suggest that *Arabidopsis* PARP2 is actively transported into the nucleus via importin-α-mediated transport. An NLS comprising - but possibly not limited to - amino acids 48-51 contributes to importin-α binding and is essential for PARP2 nuclear import.

## Acknowledgements

We thank Marcel Wiermer (University of Göttingen) for providing the pENTR/D-Topo IMPORTIN-α6 plasmid, Keehwan Kwon (J. Craig Venter Institute) for sharing plasmid pGW-nHalo and Brett Collins (University of Queensland) for sharing the pOPINE α-GFP nanobody plasmid (Addgene #49172; Kubala et al., 2010). Research in our laboratory is funded by the German Research Foundation (DFG; grant WI 3670/2-1 and Collaborative Research Centre 973 ‘Priming and Memory of Organismic Responses to Stress’) and the FU Berlin Dahlem Centre of Plant Sciences. CC is supported by the Freie Universität Berlin - China Scholarship Council (FUB-CSC) program for doctoral researchers.

## Author contributions

design of the research: LW

performance of the research: CC, RL, LW

data analysis, collection, or interpretation: CC, LW

writing the manuscript: LW

All authors have approved the submitted version of the manuscript. The authors declare that they have neither financial nor non-financial competing interests.

---

**Figure S1.**
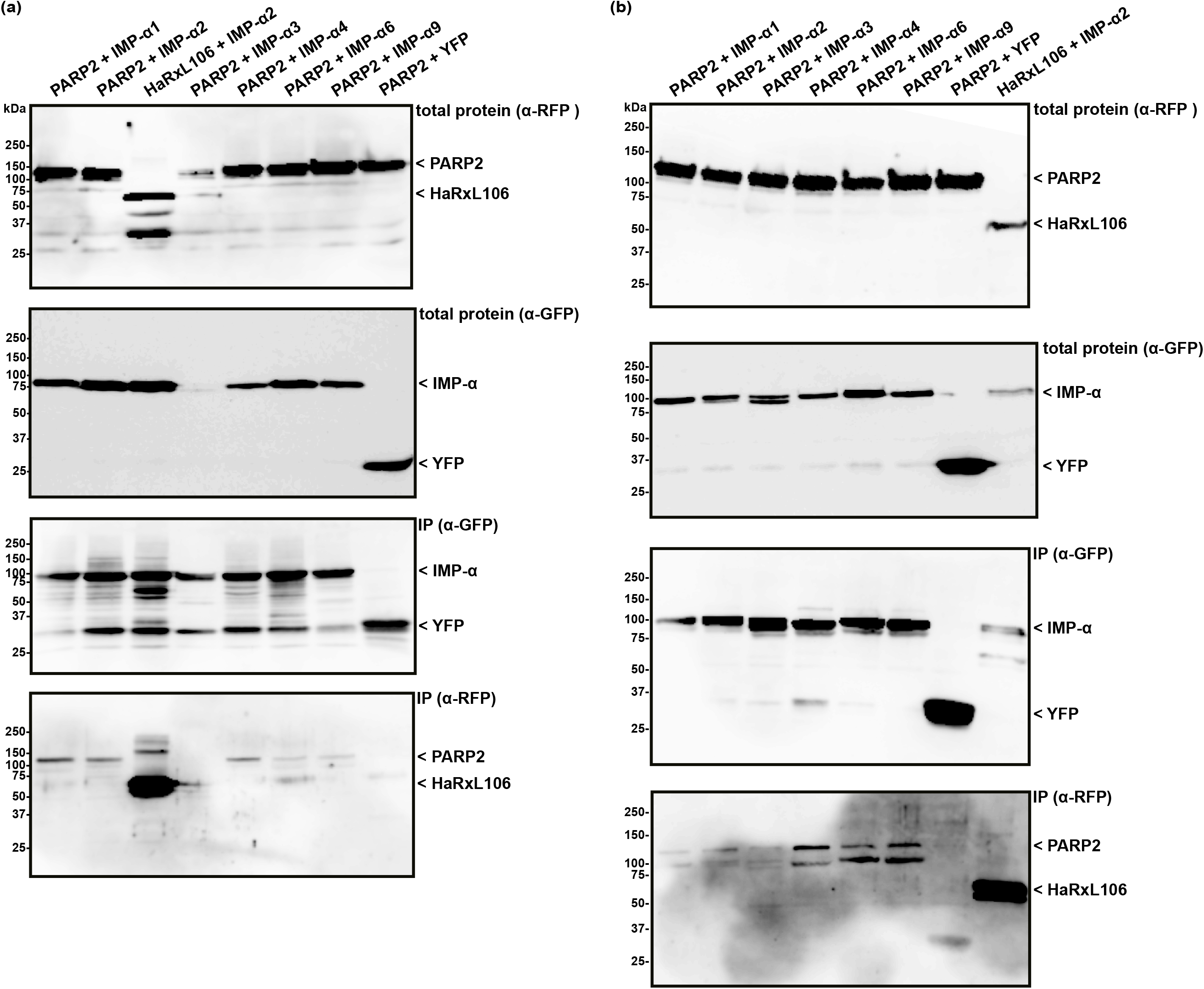
Two additional experiments of the co-immunoprecipitation experiment shown in Fig. 1b.

**Figure S2.**
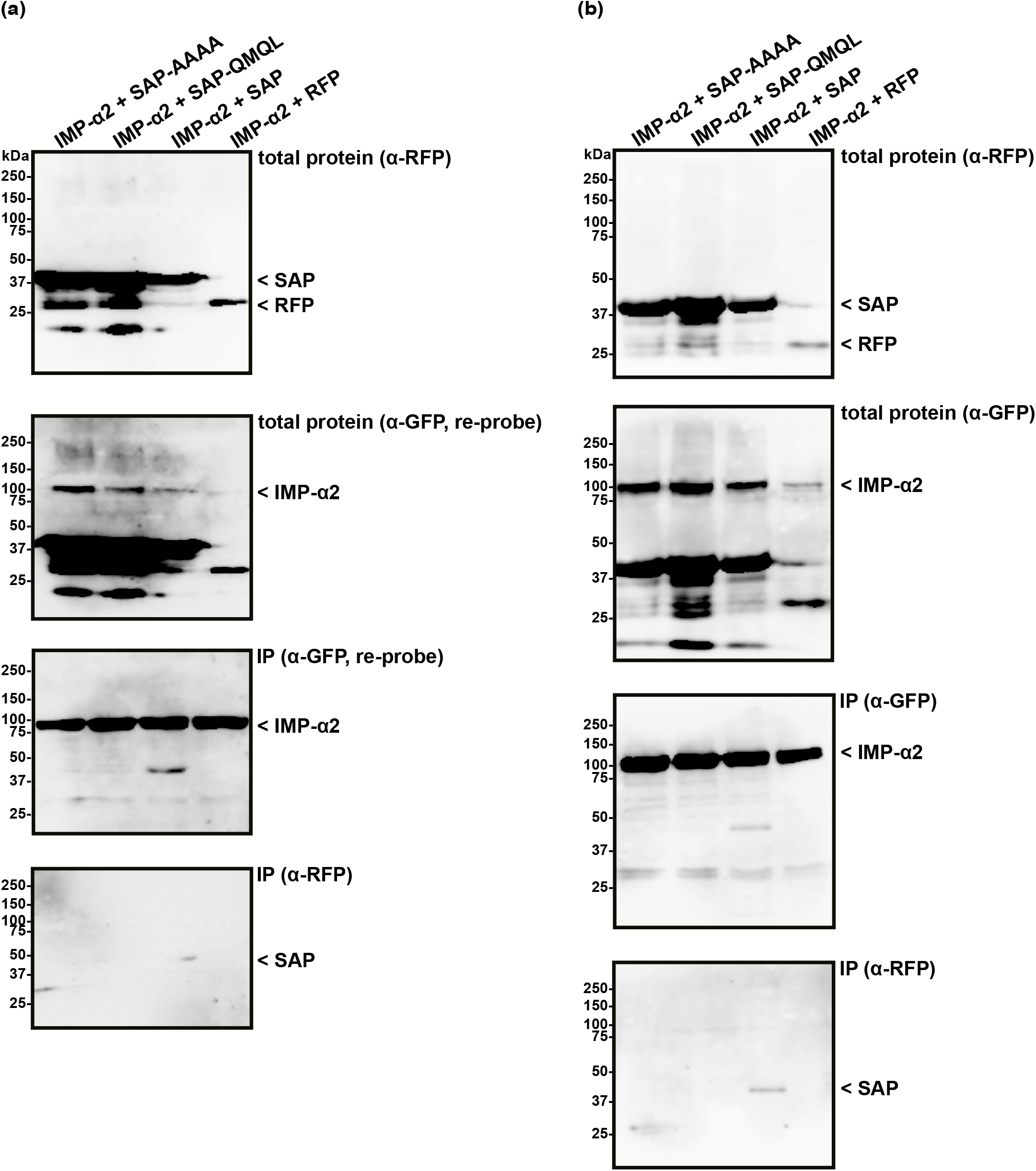
Two additional experiments of the co-immunoprecipitation experiment shown in Fig. 3c.

**Figure S3.**
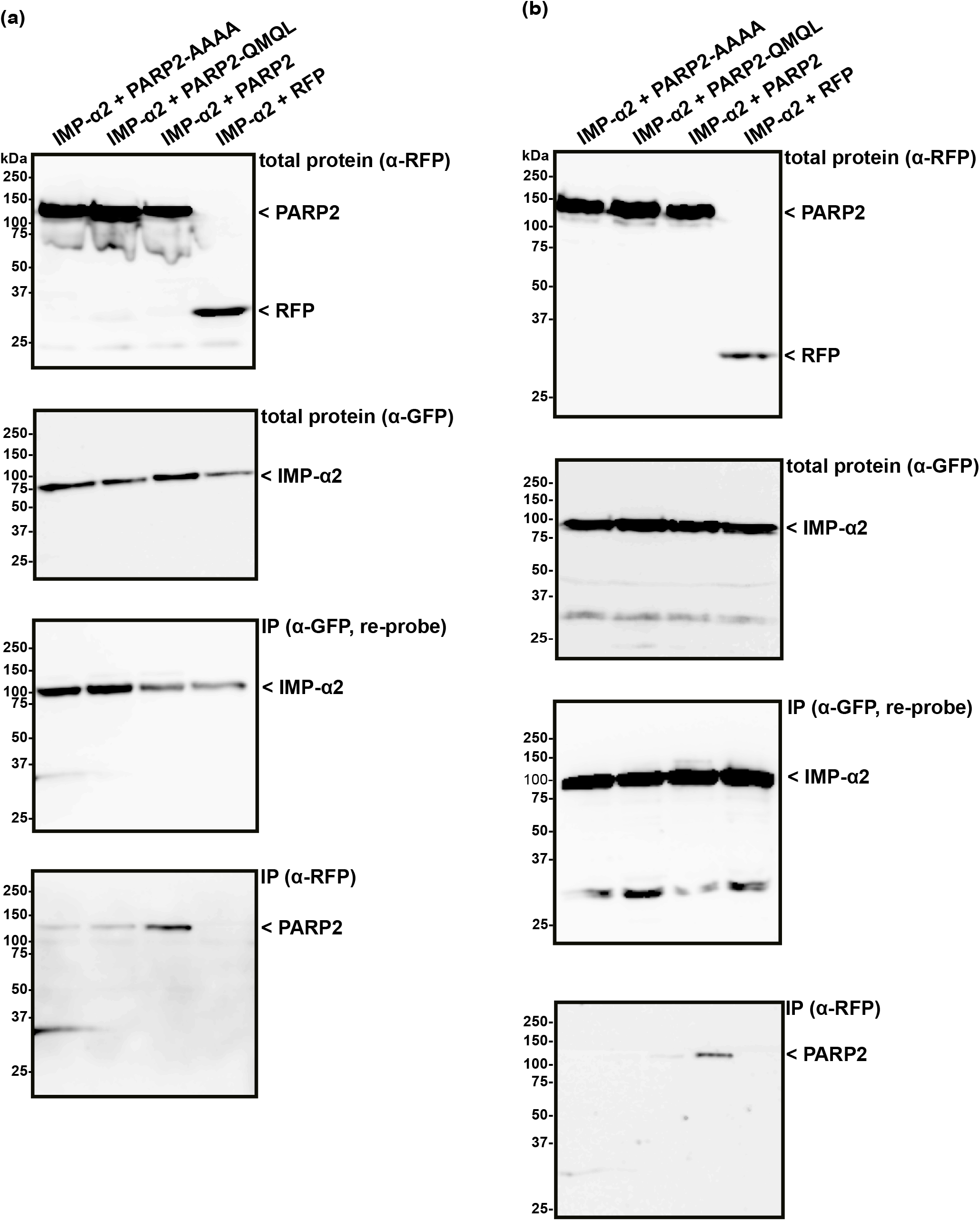
Two additional experiments of the co-immunoprecipitation experiment shown in Fig. 3d.

**Figure S4.**
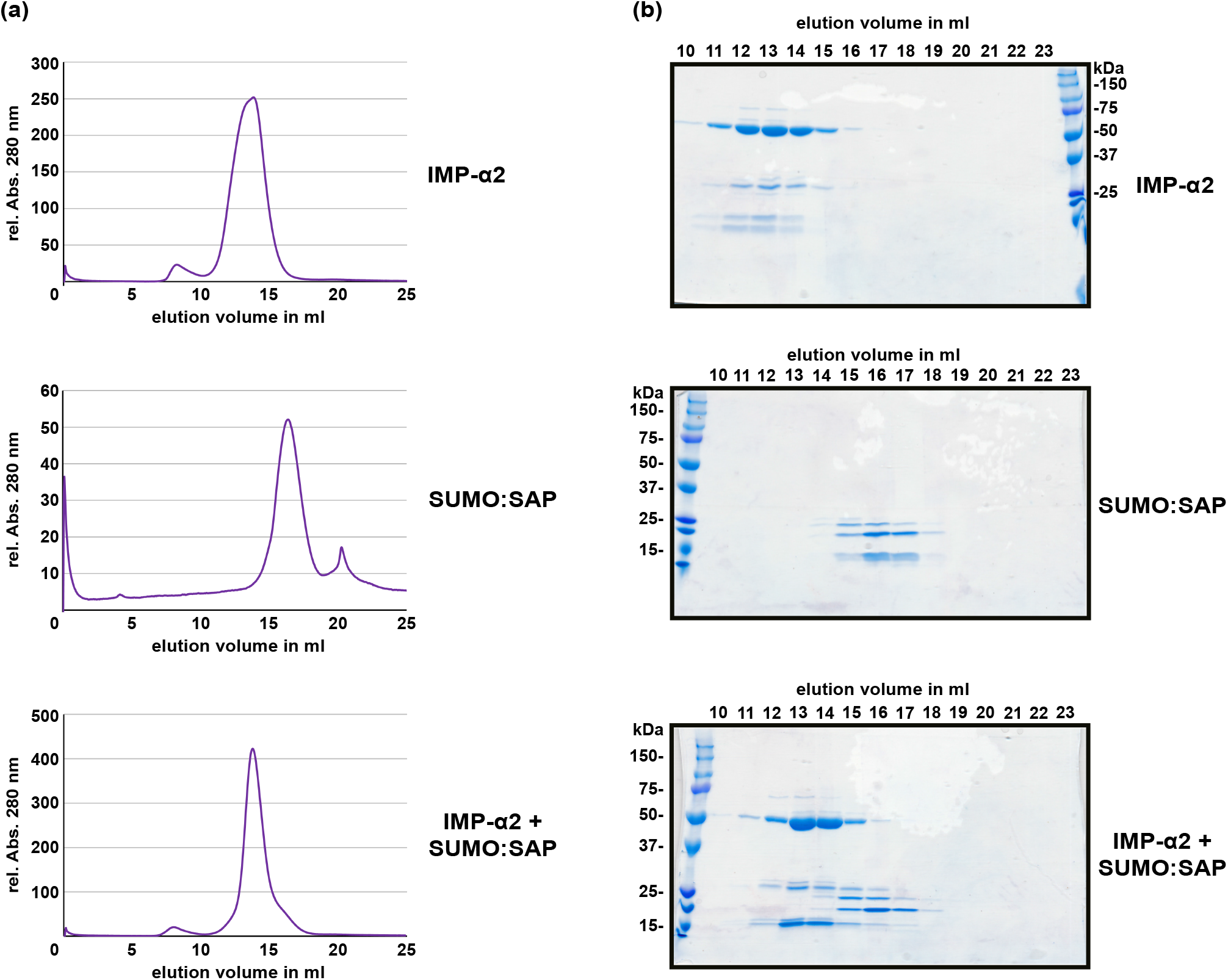
Additional experiment of the size exclusion chromatography separation shown in Fig. 4.

**Table S1.**
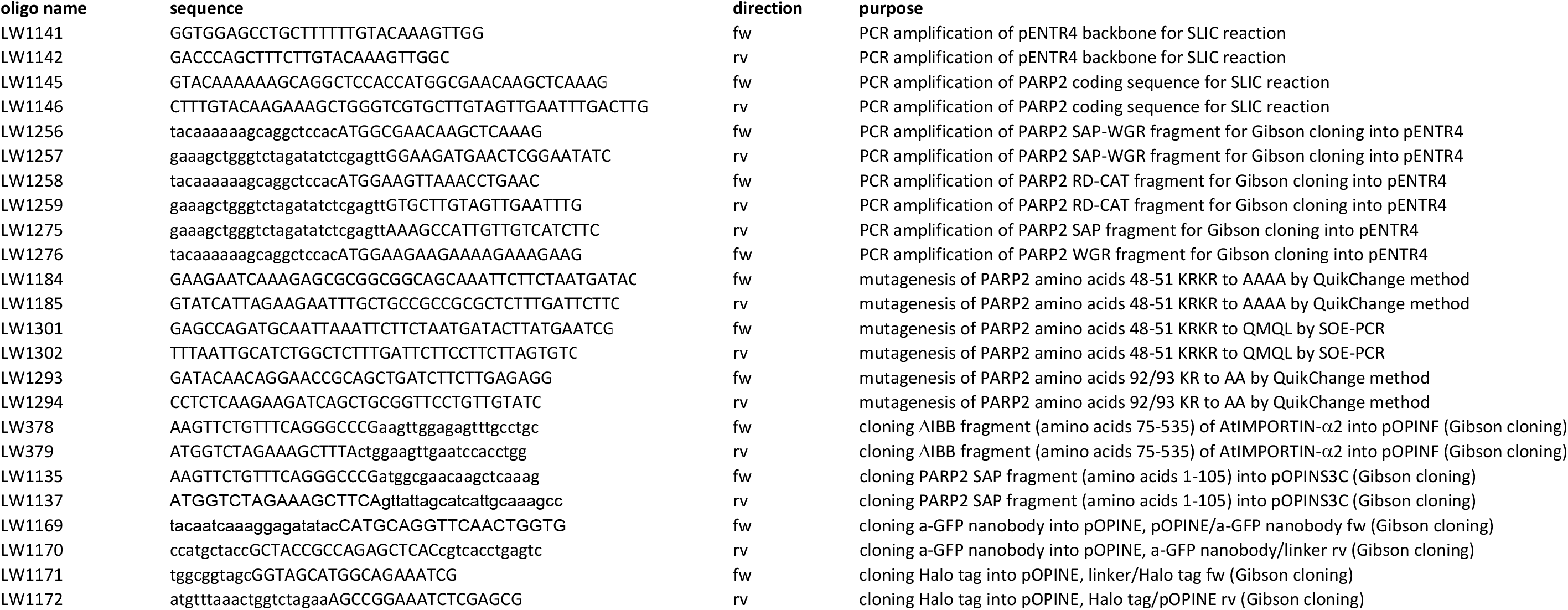
Oligo nucleotides used in this work.

